# Meiotic pairing and double strand break formation along the heteromorphic threespine stickleback sex chromosomes

**DOI:** 10.1101/2022.01.11.475862

**Authors:** Shivangi Nath, Lucille A. Welch, Mary K. Flanagan, Michael A. White

## Abstract

Double strand break repair during meiosis is normally achieved using the homologous chromosome as a repair template. Heteromorphic sex chromosomes have reduced sequence homology with one another, presenting unique challenges to the repair of double strand breaks. Our understanding of how heteromorphic sex chromosomes behave during meiosis has been largely limited to the ancient sex chromosomes of mammals, where the X and Y differ markedly in overall structure and gene content. Consequently, pairing of the X and Y chromosomes is limited to a small pseudoautosomal region. It remains unclear how more recently evolved sex chromosomes, that share considerably more sequence homology with one another, pair and repair double strand breaks during meiosis. One possibility is barriers to pairing evolve rapidly. Alternatively, recently evolved sex chromosomes may exhibit pairing and double strand break repair that more closely resembles that of their autosomal ancestors. Here we use the recently evolved X and Y chromosomes of the threespine stickleback fish *(Gasterosteus aculeatus)* to study patterns of pairing and double stranded break formation using molecular cytogenetics. We found that the sex chromosomes of threespine stickleback fish do not pair exclusively in the pseudoautosomal region. Instead, the chromosomes fully pair in a non-homologous fashion. To achieve this, the X chromosome underwent synaptic adjustment during pachytene to match the axis length of the Y chromosome. Double strand break formation and repair rate also matched that of the autosomes. Our results highlight that recently evolved sex chromosomes exhibit meiotic behavior that is reminiscent of autosomes and argues for further work to identify the homologous templates that are used to repair double strand breaks on the X and Y chromosomes.

## Introduction

Meiosis is a specialized form of cellular division process that results in the formation of haploid gametes. In order to produce functional gametes, the chromosomes have to segregate properly such that each gamete receives only one copy of each homologous chromosome pair. If chromosomes fail to pair and segregate correctly, severe meiotic errors can result (reviewed in [1]. In many species, pairing is initiated by the formation of double stranded breaks (DSBs) [2-4] that are then repaired using intact sequence from the homologous chromosome as a template. Heteromorphic sex chromosomes pairs (X/Y or Z/W) present unique challenges in terms of pairing and DSB repair during meiosis due to their overall lack of sequence homology between the chromosomes available for DSB repair [5-15].

Our understanding of how heteromorphic sex chromosomes pair and how double strand break initiation and repair occurs has largely been informed by detailed studies of the ancient X and Y chromosomes of mammals [6-10,12] and the ancient Z and W chromosome of birds [11,16]. Heteromorphic sex chromosomes share a region of complete homology (the pseudoautosomal region) that undergoes an obligate crossover each meiosis [17-20]. During meiosis, the sex chromosomes of most mammals exhibit delayed pairing and only synapse within the pseudoautosomal region, while the sex-limited, non-homologous regions remain largely unpaired [8,20-23]. Double strand breaks continue to form within the unsynapsed axial element of the X or Z chromosome [24-29], whereas they are largely suppressed on the axial element of the sex-limited chromosome [25,26]. The double strand breaks that do form within the non-homologous region exhibit delayed repair relative to the autosomes and are primarily repaired through homologous exchange with the sister chromatid or via non-homologous end joining [15]. In birds, the lack of sequence homology between much of the Z and W chromosome has resulted in error prone synapsing, with a complete failure of pairing in nearly a quarter of oocytes [11].

Although it is clear that highly degenerate sex chromosomes must navigate meiosis in a manner unique from autosomes, it is unknown how quickly these modifications are established following the evolution of sex chromosomes. Comparisons with recently evolved sex chromosomes are needed to understand how pairing and double strand break repair are altered from their autosomal ancestors. Young sex chromosomes share higher sequence homology with one another compared to older sex chromosomes that have accumulated mutations over a longer time period (reviewed in [30]). This raises the possibility that younger sex chromosomes may exhibit more extensive pairing depending on the amount of sequence and structural divergence outside of the PAR. In addition, double strand breaks may form and be repaired at rates that more closely resembles autosomes than that observed on ancient sex chromosomes. If the repair of double strand breaks occurs more often between the X and Y or Z and W through non-crossover gene conversion, these breaks could be repaired earlier in meiosis before the barrier to sister chromatid and non-homologous end joining repair is lifted [29,31].

Threespine sticklebacks (*Gasterosteus aculeatus*) are a useful species to study the behavior of recently evolved sex chromosomes during meiosis. They have an X and Y sex chromosome system which evolved only approximately 22 million years ago [32], compared to the ~180 million-year-old Y chromosome of mammals [33,34]. Crossing over between the X and Y chromosomes is suppressed across most of the chromosome pair outside of a 2.5 Mb pseudoautosomal region [35]. The sex determining region coincides with three major inversions between the X and Y, which form three separate evolutionary strata [32,36]. Despite the structural differences between the X and Y, meiotic nuclei isolated from a single population indicated the sex chromosomes may synapse fully in males along their length [37]. However, due to limited sample sizes it is unknown whether the X and Y synapse fully in a majority of meiocytes or if full pairing is mostly achieved only in the pseudoautosomal region, similar to species with ancient sex chromosomes [38]. In addition, the overall timing and rate of meiotic double strand break formation and repair is unknown between recently derived X and Y chromosomes. Synonymous sequence divergence is an order of magnitude lower in the youngest strata of the threespine stickleback X and Y chromosomes compared to the youngest strata of the human X and Y [32]. Threespine stickleback fish therefore present an ideal opportunity to explore whether double strand breaks on young sex chromosomes exhibit repair dynamics that more closely resemble that of their autosomal ancestral counterparts.

Here we use molecular cytogenetics to closely examine pairing and double strand break formation of autosomes and sex chromosomes during male meiosis. Contrary to what has been observed in species with ancient sex chromosomes, we found the more recently evolved sex chromosomes of threespine stickleback fish fully pair and repair double strand breaks at the same rate as autosomes.

## Results

### The threespine stickleback X and Y chromosomes pair non-homologously during meiosis

The Y chromosome of the threespine stickleback has undergone at least three major inversions since it diverged from the common autosome ancestor with the X chromosome [32,36]. The Y chromosome has also undergone varied degrees of sequence degeneration across much of its length, resulting in a shorter estimated chromosome size (X chromosome with pseudoautosomal region: 20.58 Mb; Y chromosome with pseudoautosomal region: 18.02 Mb; [32]). Combined, chromosome-wide synteny between the X and Y chromosomes outside of the pseudoautosomal region has been lost, suggesting the X and Y chromosomes may not fully pair during meiosis, similar to the ancient sex chromosomes of mammals [8,20-23]. To observe how the X and Y chromosomes pair, we used a well-characterized cytogenetic marker *(Idh)* to differentiate the chromosomes throughout prophase I (*Idh* is located in the middle of the X chromosome at 11.5 Mb and at the end of the Y chromosome at 18.02 Mb, opposite of the pseudoautosomal region) [32,36]. Using this marker as a probe, we found the X and Y chromosomes synapsed fully during late zygotene and pachytene, when synapsis was also complete on the autosomes (Figure 1). We found the X-linked *Idh (X-Idh)* marker is in the middle of the sex chromosome pair while the *Idh* marker on the Y chromosome *(Y-Idh)* is located further distal from the pseudoautosomal region on the synapsed chromosome pair (full synapsis was observed in 50 out of 51 nuclei). Given the large-scale structural rearrangements between the sex chromosomes [32,36], our results show synapsis of the X and Y must be proceeding in a largely non-homologous fashion. Despite this, pairing is not delayed relative to autosomes.

**Figure 1.**
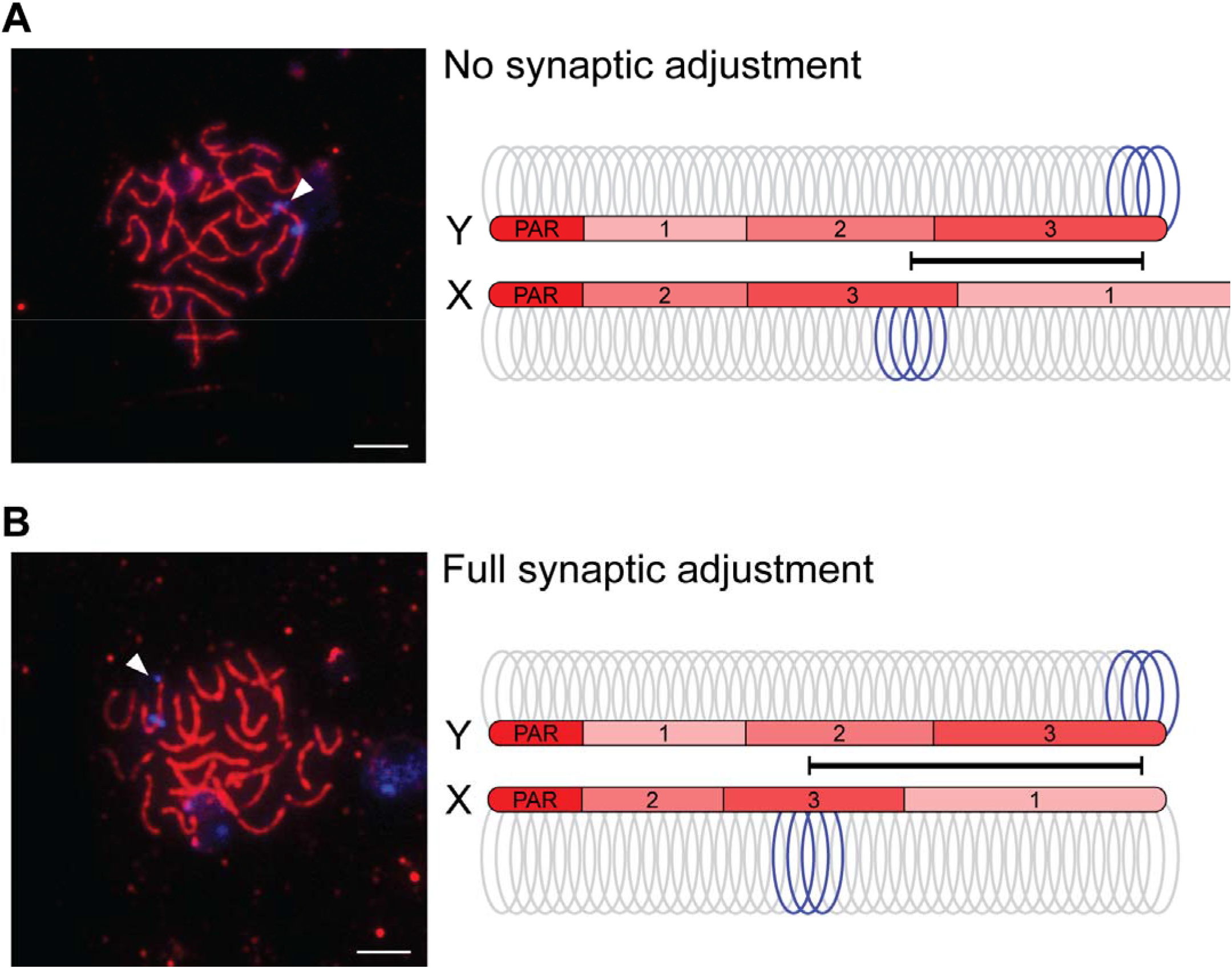
The threespine stickleback X and Y chromosomes undergo synaptic adjustment. Initially, the X chromosome synaptonemal complex axis extends beyond the length of the shorter Y chromosome (A). The sex chromosomes are marked with a fluorescent *in situ* hybridization DNA probe (*Idh*; blue). Due to chromosomal rearrangements between the X and Y, this marker is at the end of the Y chromosome, but it is located centrally within the X chromosome. Before synaptic adjustment, the axial element (SMC3; red) extends beyond the Y chromosome *Idh* marker (arrowhead). After synaptic adjustment (B), this axis no longer extends beyond the Y chromosome *Idh* marker (arrowhead). The arrangement of the three main evolutionary strata on the X and Y chromosomes (1, 2, and 3; [32]) are indicated on each axis, along with the pseudoautosomal region (PAR) where all crossing over occurs between the two chromosomes.

### The X chromosome undergoes synaptic adjustment during male meiosis

Among the fully paired X and Y chromosomes, two main conformations were identified based on the relative positions of the *Idh* markers. In one configuration, the *X-Idh* and *Y-Idh* markers were located further apart, but the Y-*Idh* marker was not at the end of the sex chromosome axis (20 out of 43 observed X/Y pairings; Figure 1A). In this configuration, the SMC3 axis extended beyond the Y chromosome *Idh* signal. In the second configuration, the *X-Idh* marker was located in the middle and the *Y-Idh* marker was located at the end of the SMC3 axis (23 out of 43 observed X/Y pairings; Figure 1B). These results are consistent with the shorter Y chromosome initially pairing with a longer X chromosome, followed by synaptic adjustment of the X to equalize the two SMC3 axes.

To test our hypothesis that synaptic adjustment was occurring between the non-homologously paired X and Y chromosomes, we first examined the overall timing of synaptic adjustment during prophase I. Synaptic adjustment does not occur until late pachytene (reviewed in [39]). Therefore, we predicted that the full adjustment configuration should be enriched in pachytene nuclei, relative to zygotene. We staged nuclei from four males to zygotene and pachytene (see methods). Consistent with synaptic adjustment, we found that the full synaptic adjustment configuration was enriched within pachytene nuclei, relative to zygotene (Figure 2A, Fisher’s exact test; *P* = 0.007).

**Figure 2.**
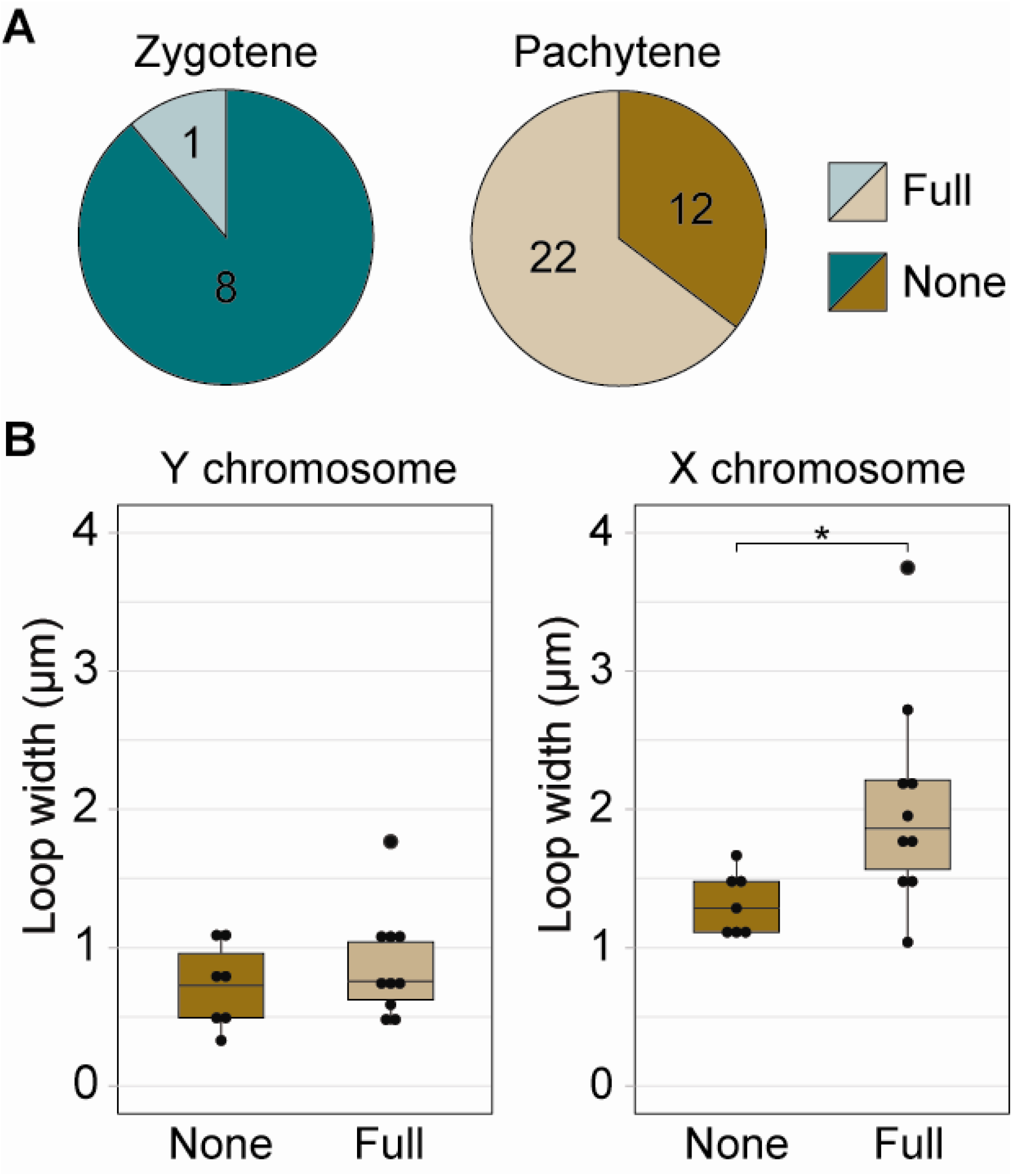
The longer X chromosome axis contracts to adjust to the shorter Y chromosome axis as meiosis proceeds. There is a greater proportion of paired X and Y chromosomes that have undergone synaptic adjustment by pachytene (A; Fisher’s exact test; *P* = 0.007). Due to the shortening axis, chromatin loops on the X chromosome are wider after full synaptic adjustment (B; Mann-Whitney U Test; *P* = 0.022). There was no change in chromatin loop width on the Y chromosome, indicating this chromosome axis is not undergoing adjustment. Chromatin loop width was measured at the *Idh* marker.

Second, we examined the size of chromatin loops organized around the SMC3 axis. The density of chromatin loops is highly conserved across taxa [39]. Given constant DNA length, shorter synaptonemal complex axes result in chromatin loops that extend further from the synaptonemal complex, whereas longer axes result in smaller loops [40]. For instance, differences in synaptonemal complex axis length between sexes of the same species results in a coordinated alteration in chromatin loop size [41,42]. If synaptic adjustment was occurring, we predicted that the chromosome undergoing adjustment would exhibit a change in loop size between the no synaptic adjustment and the full synaptic adjustment conformations. We therefore measured the width of the *X-Idh* and *Y-Idh* FISH marker signals as a proxy for DNA loop size in the no adjustment (N=7) and full adjustment (N=10) conformations among pachytene nuclei. We found that while the *Y-Idh* loop size was the same in both conformations (Figure 2B; Mann-Whitney U Test; *P* = 0.591), the *X-Idh* loop had a larger width in the full synaptic adjustment conformation (Figure 2B; Mann-Whitney U Test; *P* = 0.022). This indicates that synaptic adjustment occurs through shortening of the longer X chromosome axis. Our results are consistent with known patterns of synaptic adjustment on other chromosomes, where the longer axis shortens to match the length of the shorter axis [39,43].

### DNA double strand breaks form at a similar frequency on the autosomes and sex chromosomes

To quantify the total number of DSBs throughout the genome, we counted the total number of RAD51 foci within meiotic nuclei (Figure 3). RAD51 localizes to single-stranded DNA at DSBs and has been used to cytologically visualize sites of DSB repair [7-9,23,27,29]. We found DSBs were at the highest density within leptotene (Figure 4; N=34 nuclei; median: 0.042 RAD51 foci/Mb). As prophase proceeds, double strand breaks are repaired and the number of RAD51 foci decrease. Consistent with this, we observed a significant reduction of DSBs at each stage, decreasing as prophase proceeded (Figure 4; zygotene N=53 nuclei; median: 0.024 RAD51 foci/Mb; pachytene N=34 nuclei; median: 0.005 RAD51 foci/Mb; Mann-Whitney U Test; P < 0.001 all pairwise comparisons, Bonferroni corrected for multiple comparisons).

**Figure 3.**
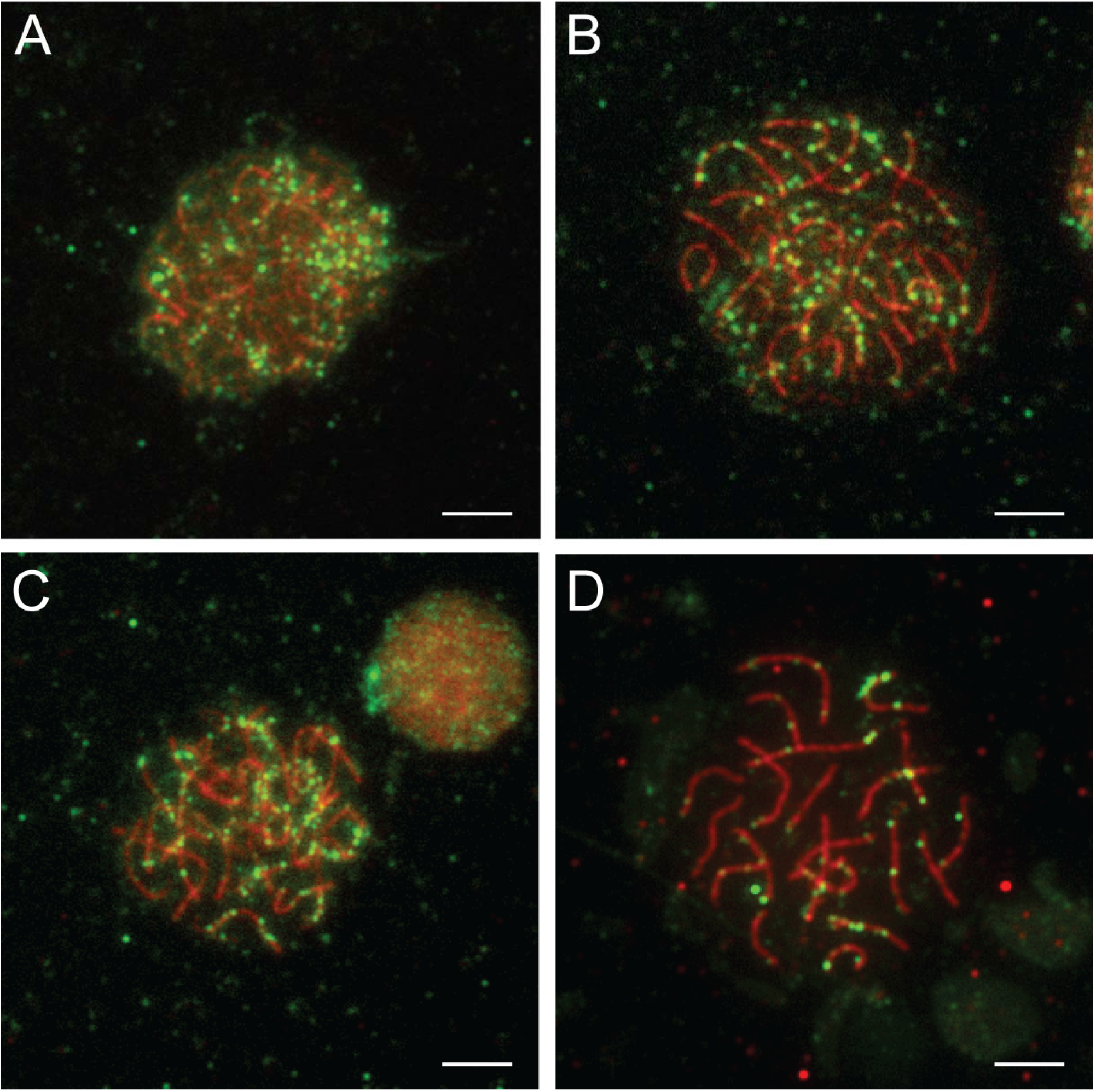
Double strand breaks form throughout the genome. The greatest number of double strand breaks occurs during leptotene (A) and decreases as meiosis proceeds through early zygotene (B), late zygotene (C), and pachytene (D). Double strand breaks are immunolabeled against RAD51 (green). The synaptonemal complex axes are immunolabeled against the axial protein, SMC3 (red).

**Figure 4.**
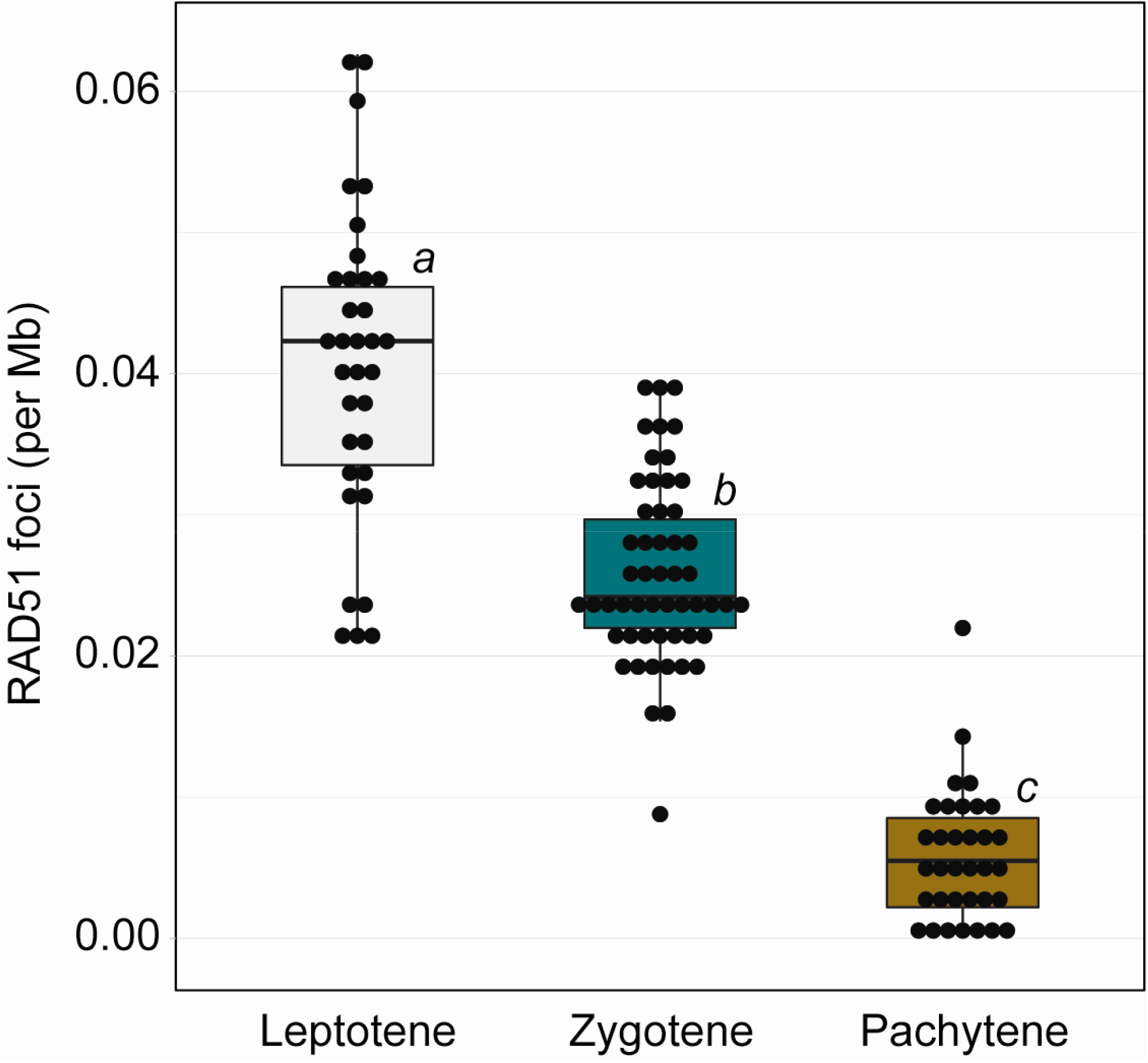
Double strand breaks throughout the genome are repaired as meiosis proceeds. The highest density of RAD51 foci throughout the genome was found during leptotene. The lowest density was observed during pachytene. Stages of meiosis significantly different from one another are indicated by *a, b,* and *c* (Mann-Whitney U Test; P < 0.05; Bonferroni corrected for multiple comparisons).

Across the degenerated sex chromosomes of mammals, DSBs are suppressed and form at a lower density, relative to the autosomes [26]. The DSBs that do form also exhibit delayed repair [8]. It is unknown whether this difference in timing is characteristic of all heteromorphic sex chromosomes, or whether less degenerated sex chromosomes may still exhibit DSB formation and repair similar to their ancestral autosome progenitors. We therefore tested whether the X and Y chromosomes of threespine stickleback fish exhibited any suppression of DSBs or whether they had counts that resembled that of autosomes. We focused on zygotene and pachytene stages when the X and Y chromosomes were fully synapsed. During zygotene, we found that the density of DSBs on the sex chromosomes was not significantly different from the density we observed on autosomes (Figure 5; N=22; Mann-Whitney U Test; P=0.327, Bonferroni corrected for multiple comparisons). Later in pachytene, the density of double strand breaks was significantly lower on the sex chromosomes, compared to the autosomes (Figure 5; N=23; Mann-Whitney U Test; P=0.003, Bonferroni corrected for multiple comparisons). Combined, our results indicate that DSBs are not suppressed on the threespine stickleback X and Y chromosomes. In fact, the formation and repair of DSBs on the sex chromosomes is coincident with the timing observed on autosomes.

**Figure 5.**
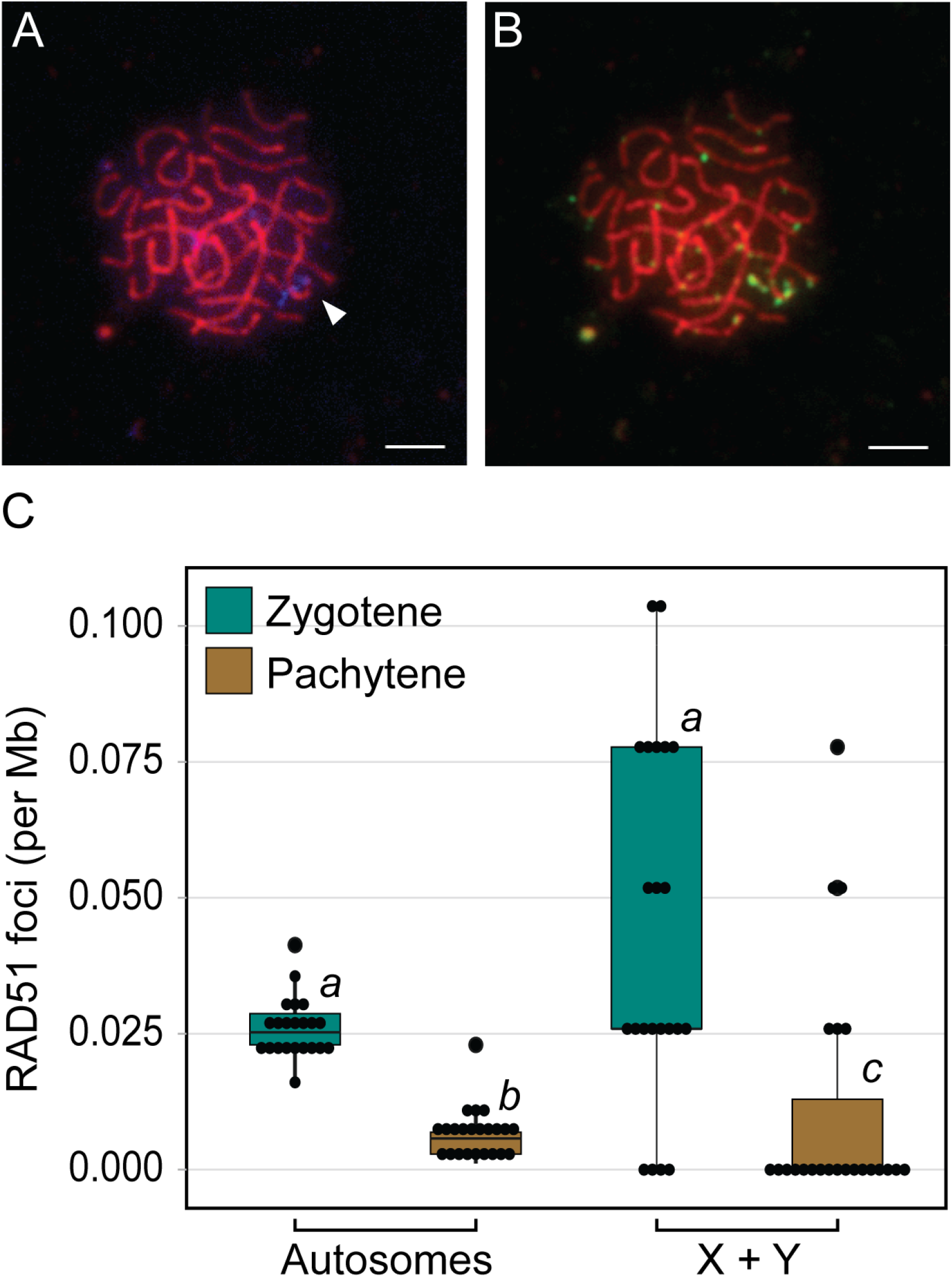
The threespine stickleback sex chromosomes have a similar density of double strand breaks as autosomes. Meiocytes at pachytene were labeled with *Idh* to distinguish the sex chromosomes (blue; arrowhead) from the autosomes (A). RAD51 foci (green) were counted on all chromosomes (B). Autosomes and sex chromosomes have an indistinguishable density of RAD51 foci at zygotene (C). The density of RAD51 foci is lower on sex chromosomes compared to autosomes at pachytene, indicating breaks may actually repair faster. Groups significantly different from one another are indicated by *a, b,* and *c* (Mann-Whitney U Test; P < 0.05; Bonferroni corrected for multiple comparisons).

### Double strand breaks on the sex chromosomes are not restricted to the pseudoautosomal region

DSBs occur at a higher density in the mammalian pseduautosomal region compared to the unsynapsed region of the X and Y chromosomes [26]. We therefore examined whether DSBs mainly occurred within the pseudoautosomal region of the threespine stickleback X and Y chromosomes or if DSBs also occurred across the remainder of the sex chromosomes where crossing over between the X and Y does not occur, despite being synapsed. We quantified the total number of DSBs in five equally sized regions across all autosomes and the sex chromosomes. The Y chromosome *Idh* marker is located at the opposite end of the chromosome from the pseudoautosomal region, which allowed us to cytologically separate the sex determining region from the pseudoautosomal region. On autosomes, we found the total number of DSBs was highest at the ends of the chromosomes (Figure 6; N=420 autosomes). On the sex chromosomes, we observed a pattern of DSB formation that closely resembled that of the autosomes. We found most of the DSBs were located at the ends of the sex chromosomes (Figure 6; N=47 X/Y chromosomes). Importantly, this includes the end of the chromosome opposite from the pseudoautosomal region, which does not undergo crossing over during meiosis. In addition, we also observed DSBs throughout the central part of the sex chromosomes that also do not undergo crossing over. Although we could not distinguish whether DSBs formed more often on the X or Y chromosome at this resolution, our results clearly indicated DSBs do occur throughout the sex determining region.

**Figure 6.**
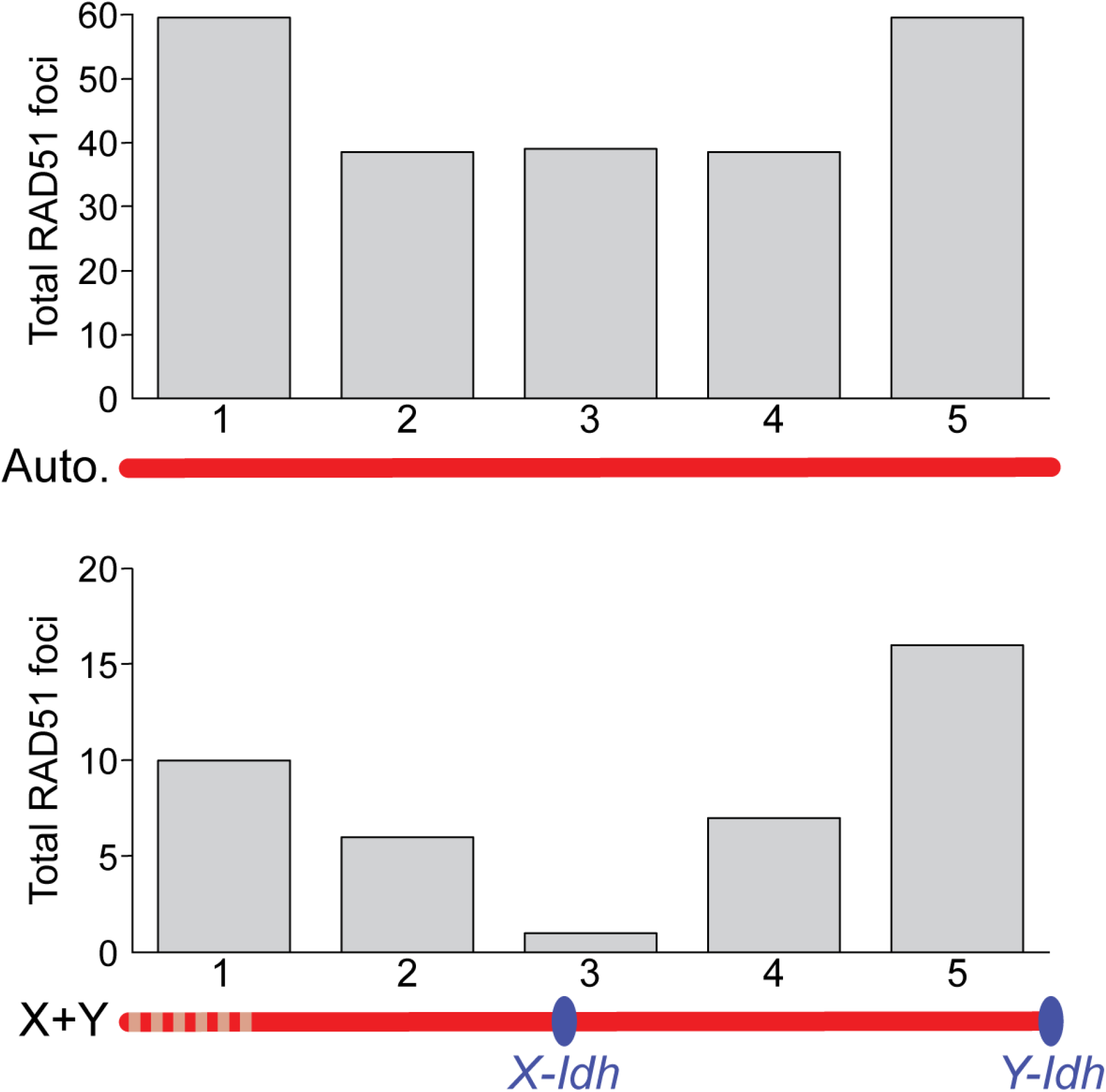
RAD51 foci occur most frequently at the terminal ends of chromosomes on both the autosomes and the sex chromosomes. Each SMC3 axis was divided into five equally sized segments and RAD51 foci were counted within each segment. The pseudoautosomal region (vertical lines) is indicated and is located at the end opposite of the terminal *Idh* marker (blue) on the paired X and Y chromosome.

### Discussion

The lack of sequence homology between heteromorphic sex chromosomes leads to pairing challenges during meiosis. The pairing of homologs requires engagement of single-stranded DNA at DSBs between homologs [3,4]. In mammals, this occurs within the pseudoautosomal region [38]. More extensive pairing outside of the pseudoautosomal region has been observed in a subset of meiocytes [22,44,45], but stable pairing is largely restricted to this short stretch of sequence homology. Unlike mammals, we observed complete pairing of the threespine stickleback heteromorphic X and Y chromosomes in nearly all meiocytes, indicating stable pairing is achieved along the entire length of the X and Y chromosomes. This pattern matches earlier work using electron microscopy that noted all chromosomes fully pair [37]. Full pairing is observed in other teleost species with more recently derived sex chromosomes. The guppy Y chromosome is recently derived [46] and fully pairs with the X chromosome [43].

However, in this species homology between the X and Y chromosomes is much higher compared to the threespine stickleback. The Y chromosome does not exhibit extensive sequence degeneration and does not contain any structural rearrangements [46-48]. In fact, rare crossover events between the X and Y are still observed in the sex determination region [49], indicating DSBs can be repaired using inter-gametolog templates. Full pairing is also observed between the Nile tilapia X and Y chromosomes [50], which harbors a sex determination region that is approximately 29% of the chromosome length [51,52]. It is unclear how the threespine stickleback X and Y chromosomes achieve full pairing during meiosis with extensive degeneration outside of the pseudoautosomal region in addition to major inversions that have occurred on the sex chromosomes [32,53]. There are no colinear regions of sequence homology between the X and Y within the sex determination region (Figure 1). In birds, full heterologous pairing occurs between the Z chromosome and the ancient, degenerated W chromosome and may be facilitated by regions of repetitive microhomology between the two gametologs [11]. Additional work will be necessary to determine if pairing of the threespine stickleback sex chromosomes is homology dependent. Knocking out *Spo11* would provide insight into whether homology-based DSB repair is required for pairing and synapsis of the sex chromosomes [3,4].

Full pairing of the threespine stickleback X and Y is achieved through synaptic adjustment of the X chromosome to match the length of the shorter Y chromosome. Synaptic adjustment of sex chromosomes has been documented in other teleost species [50,54] and the chicken [55], preventing any regions of the sex chromosomes from remaining unsynapsed. Asynapsis of sex chromosomes leads to transcriptional silencing of the chromosomes during pachytene (meiotic sex chromosome inactivation) [56,57]. Indeed, in chickens there is no evidence of meiotic sex chromosome inactivation [11], where there is synaptic adjustment. We predict that synaptic adjustment in threespine stickleback fish will also prevent meiotic sex chromosome inactivation of the sex chromosomes during pachytene. Single-cell RNA-seq will be a useful approach to test this hypothesis by characterizing transcription of the sex chromosomes relative to the autosomes at each stage of meiosis [58].

DSBs form on the threespine stickleback sex chromosomes at a density similar to autosomes. This result stands in stark contrast to the mammalian sex chromosomes, where DSBs are suppressed outside of the pseudoautosomal region [25,26]. The resolution of our experiments does not allow us to accurately place DSBs inside or outside of the pseudoautosomal region when they are near the border. However, over half of the DSBs we detected on the sex chromosomes were far from the border, at the end opposite from the pseudoautosomal region. Our results show that more recently derived sex chromosomes can accumulate DSBs with distributions that more closely resemble that of autosomes. Like the sex chromosomes, we found DSBs were enriched at the ends of autosomes. Across taxa, males often have recombination rates that are higher towards chromosome ends (reviewed in [59]). The RAD51 distribution we found matches recombination rates estimated through genetic maps in threespine stickleback fish, where males have crossover frequencies higher at chromosome ends, relative to females [60]. In humans, DSBs also occur more frequently at the end of chromosomes, suggesting DSB initiation could be the main factor that increases recombination rate within these regions [61].

The lack of sequence homology between sex chromosomes prevents repair of DSBs through homology-based mechanisms that utilize templates between gametologs. During meiosis there is a strong bias towards using the homolog as a repair template (reviewed in [62]). For highly degenerate sex chromosomes, the lack of homology between gametologs leads to a delay in repair until late pachytene [23,25,26,61] when constraints against other repair options are lifted (non-homologous end joining and homologous recombination with the sister chromatid). Components of the non-homologous end joining pathway are enriched around the mouse X and Y chromosomes during pachytene [29,63], which resembles somatic-like DNA repair, rather than meiotic-like repair that is biased towards homologous recombination with the homolog [29]. Unlike mammals, we did not find delayed DSB repair on the threespine stickleback X and Y chromosomes. The breaks repair at the same rate as DSBs located on the autosomes. This raises the intriguing possibility that DSBs on more recently derived sex chromosomes can be repaired using inter-gametolog templates when the bias against sister chromatid repair and non-homologous end joining is operating. Crossing over is completely suppressed outside of the pseudoautosomal region in the threespine stickleback fish [35,60]. Therefore, if homologous recombination is occurring between the X and Y chromosomes, DSBs must be repaired using non-crossover gene conversion pathways. This process occurs even between the ancient mammalian X and Y chromosomes at some DSBs [64-66], and may help purge deleterious mutations from the Y chromosome [67-69]. With a similar density of DSBs on the sex chromosomes as the autosomes, our results suggest gene conversion may occur at a higher rate on more recently derived sex chromosomes. Additional work will be focused on determining what DNA templates are primarily used to repair DSBs in the threespine stickleback sex determining region.

## Conclusions

Our work has revealed that recently derived sex chromosomes behave more like their ancestral autosomal progenitors during meiosis. The rates of pairing, DSB formation, and DSB repair on sex chromosomes were indistinguishable from the autosomes. Stickleback fish offer a powerful model system to explore how the meiotic behavior of sex chromosomes varies at different evolutionary scales. In addition to threespines, there are several closely related species of stickleback fish with independently derived sex chromosomes of various ages [36]. Comparative molecular cytogenetic approaches across these species will help reveal the generality of these processes over different time scales and will provide important insights into whether gene conversion between the X and Y could play a substantial role in the sequence evolution of newly evolving sex chromosomes.

## Materials and Methods

### Ethics statement

All procedures using threespine stickleback fish were approved by the University of Georgia Animal Care and Use Committee (protocol A2018 10-003-A8).

### Preparation of meiotic nuclei

Chromosome spreads were prepared using a modified protocol developed for zebrafish [70]. We targeted threespine stickleback fish that were approximately five to eight months after hatch (standard length 5.4 cm), when testes were actively undergoing meiosis [71]. Stickleback fish undergo synchronized spermatogenesis and testes of juvenile fish are enriched for similarly staged meiocytes [72,73]. Whole testes were dissected and macerated using a Dounce homogenizer in a volume of 200 μl PBS buffer. The cell suspension was centrifuged for five minutes at 200 xg and the pellet was resuspended in 200 μl of 100 mM sucrose. The cell suspension was incubated for five minutes at room temperature. 20 μl of the suspension was pipetted across a clear slide and then 100 μl of fixative (1% paraformaldehyde with 0.15% Triton-X 100) was added and the slide was left overnight in a humid chamber. The slides were washed the next day in 1X PBS three times for 15, 10, and 5 minutes and then stored at −20°C.

### Immunofluorescence of SMC3 and RAD51

We adapted an immunofluorescence protocol from Dumont et al. 2011 [74]. The spreads were permeabilized by pipetting 1.5 ml of permeabilization solution (1X PBS, 1 mM EDTA, and 1% TritonX-100) on the slide and incubating for 20 minutes at room temperature [75]. The slides were then blocked by adding 1.5 ml of 1X antibody dilution buffer and incubating for 20 minutes at room temperature. After blocking, the slides were incubated with diluted primary antibodies (1:100 anti-SMC3, ab9263; 1:40 anti-RAD51, MA5-14419) in 40 μl of 1X PBS per slide. The slides were sealed with a coverslip and rubber cement and then incubated at 4°C overnight. After the primary incubation coverslip was removed, the slides were blocked with 1X antibody dilution buffer for 20 minutes. Secondary antibodies (Goat anti-rabbit, ab150077, ab150080, and ab150079; Goat anti-mouse, ab150113) were applied at the same dilution as the primary antibodies, sealed with a coverslip and rubber cement, and incubated for two hours at 37° C. After the secondary incubation, the slides were washed in 1X PBS three times (15 minutes, 5 minutes, and 5 minutes). The slides were sealed with a coverslip and Vectashield Antifade Mounting Media (Vector Labs).

### Staging of nuclei

The spreads were staged to substages of prophase I by the following criteria: leptotene spreads were defined by small SMC3 stretches forming along chromosomes; zygotene was defined by long SMC3 stretches covering entire chromosomes undergoing synapsis, although not all homologs were fully paired (the number of chromosomes at this stage ranged from 42 unpaired homologs to two unpaired homologs); and pachytene was characterized as all homologs being paired, with a total diploid chromosome count of 21.

### Fluorescence *in situ* hybridization

The X and Y chromosomes were distinguished using the previously characterized sex-linked bacterial artificial chromosomes (BAC) clone, 101E08 (*Idh)* from the CHORI-213 library [53,76]. The BAC containing *Idh* was extracted from 200 ml bacterial cultures using the NucleoBond Xtra Midi kit (Takara Bio). The fluorescence *in situ* hybridization (FISH) probes were made using a Vysis Nick Translation Kit (Abbott Molecular) following the manufacturer’s instructions, except for the addition of two micrograms of input DNA for each reaction, rather than the one microgram suggested. The reagents were mixed and incubated for 16 hours at 15°C, followed by inactivation for 15 minutes at 70°C. After nick translation, 10ul of each probe was precipitated with 1 μl of salmon sperm DNA (ThermoFisher Scientific), 30.3 μl of 100% ethanol and 1.1 μl of 3M sodium acetate, over 15 minutes at room temperature in the dark. The probes were then centrifuged for 30 minutes at 4°C at 12,000 rpm. The supernatant was discarded, and the pellet was left to dry at room temperature for 15 minutes. The pellet was then reconstituted in 2 μl of TE (pH 8.0) and 8 μl of hybridization buffer (5 mL 100% formamide, 1 mL 20x SSC pH 7.0, 2 mL 50% dextran sulfate). The FISH probes were hybridized to the same slides used for immunofluorescence. The slides were incubated with 2X SSC at 75° C for five minutes and then treated with denaturing solution (2X SSC, 70% formamide) for two minutes. The slides were then passed through an ethanol series (70%, 85% and 100%) for two minutes each and dried in a slanted position for 10 minutes. 10 μl of the precipitated and reconstituted probe was put on the slide and sealed with a coverslip and rubber cement. The slides were incubated overnight at 37° C and washed the next day three times at 45°C in 2x SSC/50% formamide (pH 7.0), followed by three washes at 45°C in 2X SSC. Each wash was five minutes each. After drying, the slides were mounted with Vectashield Antifade Mounting Media (Vector Laboratories). All slides were imaged using a Leica DM6000 B upright microscope at 63x magnification with DAPI, TRITC, Alexa633 and FITC filter sets. All images were captured using a Hamamatsu ORCA-ER digital camera.

### Quantification of synaptic adjustment

Synaptic adjustment was only measured in nuclei where the sex chromosomes were fully paired and there were two clear *Idh* foci, one from the X chromosome and one from the Y chromosome. Sex chromosomes from pachytene and zygotene stages from four different males were categorized into two different conformations. In one conformation, the X chromosome *Idh* was located at the center of the paired chromosomes axis and the Y chromosome *Idh* was at an intermediate position between the center *Idh* and the chromosome end. In the second conformation, the X chromosome *Idh* was located centrally and the Y chromosome *Idh* was at the very end of the paired chromosome axis (Figure 1).

We examined whether the size of chromatin loops extending from the synaptonemal complex axis differed among the conformations by measuring the total width of the X chromosome and Y chromosome *Idh* marker signals extending from SMC3 using ImageJ (https://imagej.nih.gov/ij/). The freehand tool was used to draw a line connecting the two ends of each *Idh* probe that extended from the chromosome axis. The length of each line was measured in pixels and converted to microns by multiplying by a factor of 0.13 microns/pixel.

For both conformations, the number of RAD51 foci was also counted. To analyze the distribution of RAD51 foci across the sex chromosomes, the length of each chromosome was measured using ImageJ and then divided into five equal parts based on the total length of the SMC3 axis. The number of DSBs were then recorded in each interval. A total of 420 autosomes and 47 sex chromosomes from spreads in zygotene/pachytene were counted.

### Double strand break counting and normalization

For each meiotic nucleus, the total number of RAD51 foci were counted on the autosomes and the sex chromosomes. The counts were repeated three times and the average was used. The counting was conducted blinded in respect to the substage of prophase. The density of DSBs per Mb was calculating by dividing the genome-wide counts by the diploid genome size of 910.06 Mb, while the number of DSBs on the sex chromosomes was divided by 38.60 Mb (the total size of the X chromosome and Y chromosome combined).

## Acknowledgements

The authors thank Shaugnessy McCann, Jackie Cory, Jim Cory, Damean McCann, and Bill Frothingham for assistance collecting threespine stickleback fish. Kelly Dawe, Beth Dumont and Kyle Swentowsky provided assistance in troubleshooting and useful discussions of our results. Brittany Dorsey helped count RAD51 foci.

## Competing Interests

The authors do not have any competing interests to declare.

## Notes

### Competing Interest Statement

The authors have declared no competing interest.

